# The chemoreceptor controlling the Wsp-like transduction pathway in *Halomonas titanicae* KHS3 binds purine derivatives

**DOI:** 10.1101/2024.04.09.588698

**Authors:** Fernando E. Ramos Ricciuti, M. Karina Herrera Seitz, Ana F. Gasperotti, Alexandra Boyko, Kirsten Jung, Marco Bellinzoni, María-Natalia Lisa, Claudia A. Studdert

## Abstract

The chemosensory pathway *Ht*Che2 from the marine bacterium *Halomonas titanicae* KHS3 controls the activity of a diguanylate cyclase. Constitutive activation of the pathway results in colony morphology alterations and increased ability to form biofilm. Such characteristics resemble the behaviour of the Wsp pathway of *Pseudomonas*. In this work we investigate the specificity of Htc10, the only chemoreceptor coded within the *Ht*Che2 gene cluster. Thermal shift analyses performed with the Htc10 ligand-binding domain led to the identification of purine derivatives as ligands. This ligand-binding domain was crystallized in the presence of guanine or hypoxanthine and its structure was solved by X-ray protein crystallography. The sensor domain adopts a double-cache folding, with ligands bound to the membrane-distal pocket. A high-resolution structure of the occupied guanine-binding pocket allowed the identification of the involved residues. These residues were validated by site directed mutagenesis and thermal shift or isothermal calorimetry analyses of the protein variants. The dissociation constants for guanine or hypoxanthine of the intact domain were in the low micromolar range. To our knowledge, this is the first description of binding specificity for a chemoreceptor that controls the activity of an associated diguanylate cyclase, and opens the way for dynamic studies of the signalling behaviour of this kind of sensory complex. A comparison between Htc10 and the functionally equivalent WspA receptor from *Pseudomonas* revealed no significant sequence similarities. In contrast, highly conserved Htc10-like receptors were found in distant bacteria carrying *Ht*Che2-like clusters.

## Introduction

Bacteria can sense their chemical surroundings and make decisions in response. Bacterial chemosensing represents one of the most studied signal transduction systems at the molecular level, and yet poses enigma and unresolved questions. Most of what we know about the subject originates in studies from the sixties about chemotaxis in enteric bacteria that were later extended to Bacteria and Archaea showing an amazingly strong conservation with delicate variations [1,2].

*Halomonas titanicae* KHS3 is an environmental bacterium isolated from the Argentine Sea. Its genome sequence revealed the presence of two chemotaxis-related gene clusters [3]. Cluster Che1 (*Ht*Che1) resembles the Che cluster from *Escherichia coli* and controls the chemotaxis behaviour [4]. Cluster Che2 (*Ht*Che2), instead, contains a diguanylate cyclase gene encoding a protein with N-terminal phosphorylatable domains (*Ht*DGC, locus RO22_21180), resembling the Wsp pathway of *Pseudomonas* [3] (see below). Moreover, gene cluster *Ht*Che2 seems to be functionally equivalent to the Wsp cluster as a deletion mutant in the methylesterase gene *cheB2*, predicted to cause a constitutive activation of the pathway, resulted in a wrinkly colony phenotype and in increased biofilm formation ability [4].

Most chemotaxis receptors consist of homodimeric transmembrane proteins with an extracellular domain for ligand binding and an intracellular cytoplasmic domain [5]. The latter interacts with other chemoreceptors, with a coupling protein and a histidine kinase. The kinase activity is responsible for transmitting the information from chemoreceptors to the flagellar motor via another protein whose interaction depends on its phosphorylation state. Two additional enzymes that methylate or demethylate the cytoplasmic domain of chemoreceptors within a specific region are responsible for the adaptation to persistent stimuli, counteracting the activity change generated by ligands. In all, the combination of stimuli-triggered changes and the adaptation ability allows bacteria to respond to gradients along several orders of magnitude [1]. With little variations, most signal transduction systems that are responsible for the chemotactic behaviour of Bacteria or Archaea adjust to such general model. However, very similar signal transduction systems have evolved to control alternative functions other than chemotaxis [2]. One of such systems has been described to control the occurrence of the so-called <u>w</u>rinkly <u>s</u>preader <u>p</u>henotype (Wsp) in bacteria from the genus *Pseudomonas*through the phosphorylation-mediated control of an associated diguanylate cyclase [6,7]. The resulting raise in c-di-GMP levels generated by enzymatic activation results in an increased ability to form biofilm, and mutants with a constitutively activated pathway develop the rough and wrinkly colonies that gave the name to the characteristic phenotype.

Most of the 25 chemoreceptor genes coded in *Halomonas titanicae* KHS3 genome belong to the same length-class and are supposed to feed information into the chemotaxis pathway [3]. Contrastingly, the only chemoreceptor coded within the cluster *Ht*Che2, Htc10 (locus RO22_21155), possesses a longer signalling hairpin, suggesting that it assembles with the rest of the cluster *Ht*Che2-coded proteins and signals independently from the chemotaxis pathway [3,8].

Like WspA, the chemoreceptor from the *Pseudomonas* Wsp pathway, Htc10 contains a periplasmic domain flanked by two transmembrane segments, but the predicted structure of the ligand-binding domain (LBD) is of the double-cache type [3], contrasting with the 4-helix bundle structure of WspA LBD. Up to now, there are no reports about ligand-binding abilities for the periplasmic domain of chemoreceptors coded within Wsp-like gene clusters, and the current view about WspA signalling focuses in some sort of surface-sensing ability that functions independently of its LBD [9].

In this work, we have recombinantly produced the LBD of Htc10 (Htc10-LBD), with the aim to identify its specific ligand/s. Thermal shift analyses against a library of compounds identified purine-derivatives guanine and hypoxanthine as putative ligands, which were confirmed by isothermal titration calorimetry. Also, we solved a high-resolution crystal structure of Htc10-LBD in complex with guanine, which allowed identifying the residues involved in ligand binding. The involvement of such residues in purine binding was validated by mutational studies.

Up to our knowledge, this is the first report for specific ligands of a chemotaxis-related receptor presumably controlling the activity of a diguanylate cyclase. This discovery might pave the way for dynamic signalling studies of these relatively unexplored pathways.

## Results and Discussion

### Thermal shift assays identify purine derivatives as Htc10-LBD ligands

To identify the periplasmic domain of Htc10, the amino acid sequence of the predicted protein was analyzed using DeepTMHMM [10]. Two transmembrane segments were clearly recognized consisting of residues 14-32 and 307-327. The coding sequence for amino acids 32-309 (hereafter Htc10-LBD) was cloned into plasmid pET28a, in order to produce the recombinant protein fused to a C-terminal 6-His tag. Attempts to express Htc10-LBD in BL21 cells were unsuccessful. However, the co-transformation of pET28a-Htc10-LBD with the chaperone-coding plasmid pKJE7 [11] allowed the production of high amounts of soluble Htc10-LBD even in the absence of arabinose, the inducer for chaperones expression. Htc10-LBD was purified by affinity chromatography and size exclusion chromatography.

Htc10-LBD was then used in thermal shift assays to screen a library of 386 compounds (Biolog plates PM1 to PM5) including carbon, nitrogen, phosphor and sulfur sources as well as nutritional supplements. In the absence of ligands, the melting temperature (*T*_m_) of Htc10-LBD was 54.2 °C (Figure 1A). The presence of compounds containing nitrogenous bases changed the *T*_m_ by more than 2 °C (Table 1). When such compounds were tested individually at a concentration of 10 mM, the purine bases guanine and hypoxanthine produced an increase in *T*_m_ of 6 °C and 16 °C, respectively (Table 1), indicating their ability to stabilize Htc10-LBD. Besides, when tested at different concentrations, these two compounds led to a dose-dependent behavior of Δ*T*_m_, which was used to calculate dissociation constants (*K*_d_) as previously described [12] (Figure 1B). Estimations obtained with this method provided *K*_d_ values in the low micromolar range (1.6 µM for hypoxanthine and 2.3 µM for guanine). These values fall well within the range of dissociation constants reported for double-cache chemoreceptors, known to bind a variety of ligands (amino acids, taurine, citrate, polyamines, histamine) [13].

**Figure 1.**
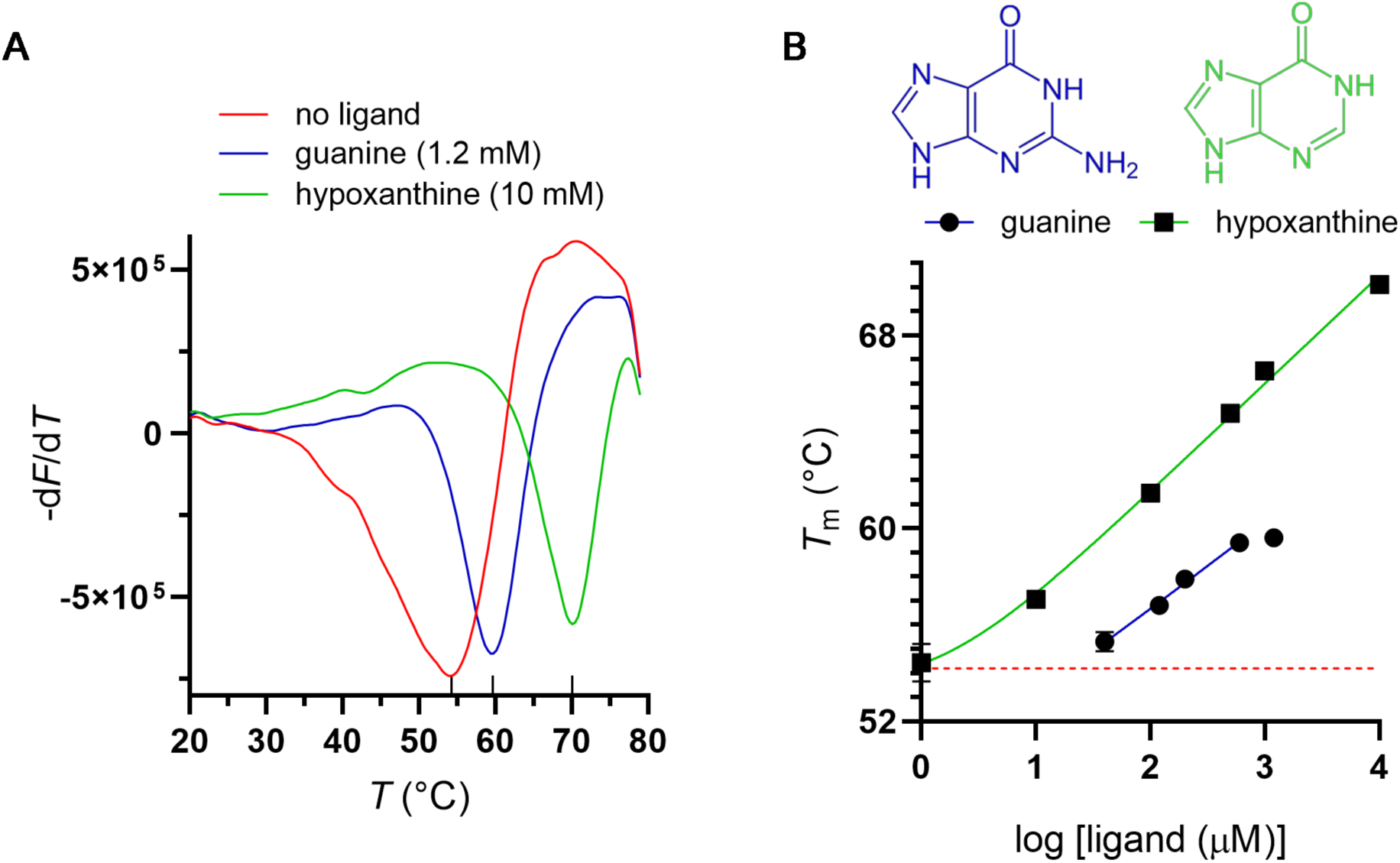
*K*_d_ determination by thermal shift assays. **(A)** Experiments were performed in the absence of ligand (red) or in the presence of 10 mM hypoxanthine (green) or 1.2 mM guanine (blue). The first derivatives of the corresponding fluorescence curves are shown; the minimum indicates the melting temperature in each case. **(B)** *T*_m_ was determined at different hypoxanthine or guanine concentrations and *K*_d_ was calculated by fitting the data to the equation *T*_m_ = *T*_m0_ + A. log (1+ 10^log[ligand]^/*K*_d_), where *T*_m0_ represents the melting temperature in the absence of ligand [12]. Chemical structures of the assayed compounds are shown.

**Table 1.** *T*_m_ shift observed for Htc10-LBD in the presence of different compounds. The difference (Δ*T*_m_) between *T*_m_ values in the presence versus in the absence of ligand is shown from representative experiments. Initial screening of compounds from Biolog plates was performed as described in Methods. ND: not determined.

| Compound | $\Delta T_m$ (°C) | |
| --- | --- | --- |
|  | Biolog plates | 10 mM solution |
| Inosine | 3.0 | 3.0 |
| **Guanine** | **6.0** | **6.0** |
| Guanosine | 2.5 | 0.5 |
| Thymine | 2.0 | 1.7 |
| Thymidine | 2.0 | 0.0 |
| Uracil | 3.0 | 3.0 |
| Xanthine | 2.5 | -8.0 |
| Guanosine 2-monophosphate | 2.5 | ND |
| **Hypoxanthine** | **8.0** | **16.0** |
| 2-deoxy-inosine | 6.5 | ND |

### Htc10-LBD is a dimer-forming double-cache domain

It has been predicted that Htc10-LBD would adopt the topology of a double-cache domain [3]. In order to test this hypothesis, we solved the crystal structure of Htc10-LBD. Due to the low sequence identity (17%) of Htc10-LBD with the LBD of *Pseudomonas fluorescens* chemoreceptor CtaA (CtaA-LBD; PDB code 6PY5 [14]), its closest match in the PDB retrieved by the HHpred server [15], we prepared seleno-methionine (Se-Met)-labeled Htc10-LBD (Se- Met-Htc10-LBD) for experimental phasing.

Se-Met-Htc10-LBD crystallized in the presence of guanine or hypoxanthine. Se-Met-Htc10- LBD_GUA_ and Se-Met-Htc10-LBD_HYP_ crystals were isomorphous, belonged to the space group H32 and diffracted to 3.6 Å (Table 2). The crystal structures of Se-Met-Htc10-LBD_GUA_ and Se- Met-Htc10-LBD_HYP_ contain a single protein molecule in the asymmetric unit, with the model of the polypeptide comprising Htc10 residues 48-230 and 264-296, while the electron density vanishes for the most N- and C-terminal residues as well as for the 231-263 segment, possibly reflecting the high flexibility of such regions. The RMSD between the protein chains in the two structures is 0.6 Å for 211 aligned alpha carbons.

**Table 2.** X-ray diffraction data collection and refinement statistics.

|  | Se-Met-Htc10-LBD<sub>GUA</sub><br>(PDB 9B9S) | Se-Met-Htc10-LBD<sub>HYP</sub><br>(PDB 9B9X) | Htc10-LBD<sub>GUA</sub><br>(PDB 9BA3) |
| --- | --- | --- | --- |
| **Data collection** |  |  |  |
| Space group | H32 | H32 | P2<sub>1</sub>2<sub>1</sub>2<sub>1</sub> |
| Cell dimensions |  |  |  |
| *a*, *b*, *c* (Å) | 178.54 178.54 89.38 | 175.86 175.86 88.61 | 89.35 104.50 145.61 |
| $\alpha$, $\beta$, $\gamma$ (°) | 90 90 120 | 90 90 120 | 90 90 90 |
| Resolution range | 77.38-3.6 (3.94-3.60)* | 76.59-3.63 (3.98-3.63) | 89.35-2.1 (2.14-2.10) |
| *R*<sub>merge</sub> (within I+/I-) | 0.081 (0.532) | 0.203 (1.080) | 0.066 (0.892) |
| *R*<sub>merge</sub> (all I+ and I-) | 0.124 (0.836) | 0.214 (1.122) | 0.093 (1.310) |
| *R*<sub>pim</sub> (within I+/I-) | 0.043 (0.282) | 0.091 (0.485) | 0.033 (0.434) |
| *R*<sub>pim</sub> (all I+ and I-) | 0.046 (0.307) | 0.066 (0.345) | 0.032 (0.441) |
| *I*/σ*I* | 8.2 (1.6) | 10.3 (2.5) | 10.2 (1.1) |
| CC (1/2) | 0.998 (0.828) | 0.996 (0.802) | 1.000 (0.759) |
| Completeness (%) | 100 (100) | 99.9 (99.9) | 100 (100) |
| Multiplicity | 8.1 (8.4) | 11.3 (11.5) | 9.4 (9.7) |
| **Refinement** |  |  |  |
| Resolution (Å) | 51.54-3.6 (4.54-3.6) | 57.75-3.63 (4.57 -3.63) | 84.9-2.1 (2.12-2.1) |
| No. reflections | 6437 (3021) | 6038 (2834) | 80127 (2589) |
| *R*<sub>work</sub>/*R*<sub>free</sub> | 0.25/0.29 | 0.24/0.29 | 0.21/0.24 |
| Protein residues | 216 | 214 | 947 |
| Ligand molecules | 1 | 1 | 6 |
| No. atoms |  |  |  |
| Protein | 1714 | 1697 | 7555 |
| Ligand | 11 | 10 | 66 |
| Water | 0 | 0 | 570 |
| Wilson B-factor | 124.54 | 109.69 | 37.79 |
| B-factors (Å<sup>2</sup>) |  |  |  |
| Protein | 114.99 | 117.00 | 48.26 |
| Ligand | 122.44 | 100.48 | 40.74 |
| Water | - | - | 46.42 |
| R.m.s. deviations |  |  |  |
| Bond lengths (Å) | 0.003 | 0.002 | 0.004 |
| Bond angles (°) | 0.57 | 0.54 | 0.66 |
| Ramachandran |  |  |  |
| Favored (%) | 97.6 | 98.1 | 98.61 |
| Allowed (%) | 2.4 | 1.9 | 1.39 |
| Outliers (%) | 0 | 0 | 0 |

\*One protein crystal was employed for structure determination in each case. Values in parentheses are for highest-resolution shell.

Se-Met-Htc10-LBD adopts the topology of a double-cache domain (Figure 2), with a RMSD of 3.0 Å for 186 residues aligned with the crystallographic model of CtaA-LBD (PDB code 6PY5 [14]) by the Dali server [16]. Like other members of this structural family of proteins [17], Htc10- LBD presents a main axis, that would be arranged perpendicular to the plasma membrane, composed by helices α1 and α2, on which the membrane-distal and the membrane-proximal pockets sit. The distal pocket consists of a β-sheet made by six β-strands (β1-6) and two α- helices, one (α3) at the N-terminus of the domain and another (α4) facing the β-sheet. The proximal pocket is made of α5 and a central nucleus of five β-strands (β7-11) (Figure 2B). Notably, a molecule of Se-Met-Htc10-LBD interacts with a nearby crystallographic symmetry mate in such a way that a dimer with C2 symmetry is formed that resembles the quaternary structure of LBDs in double-cache chemoreceptors [18] (Figure 2A). The interface area in Se- Met-Htc10-LBD dimers is *ca.* 1000 Å^2^ and the dimeric interaction is stabilized by hydrogen bonds and salt bridges among residues comprised in segment 70-107.

**Figure 2.**
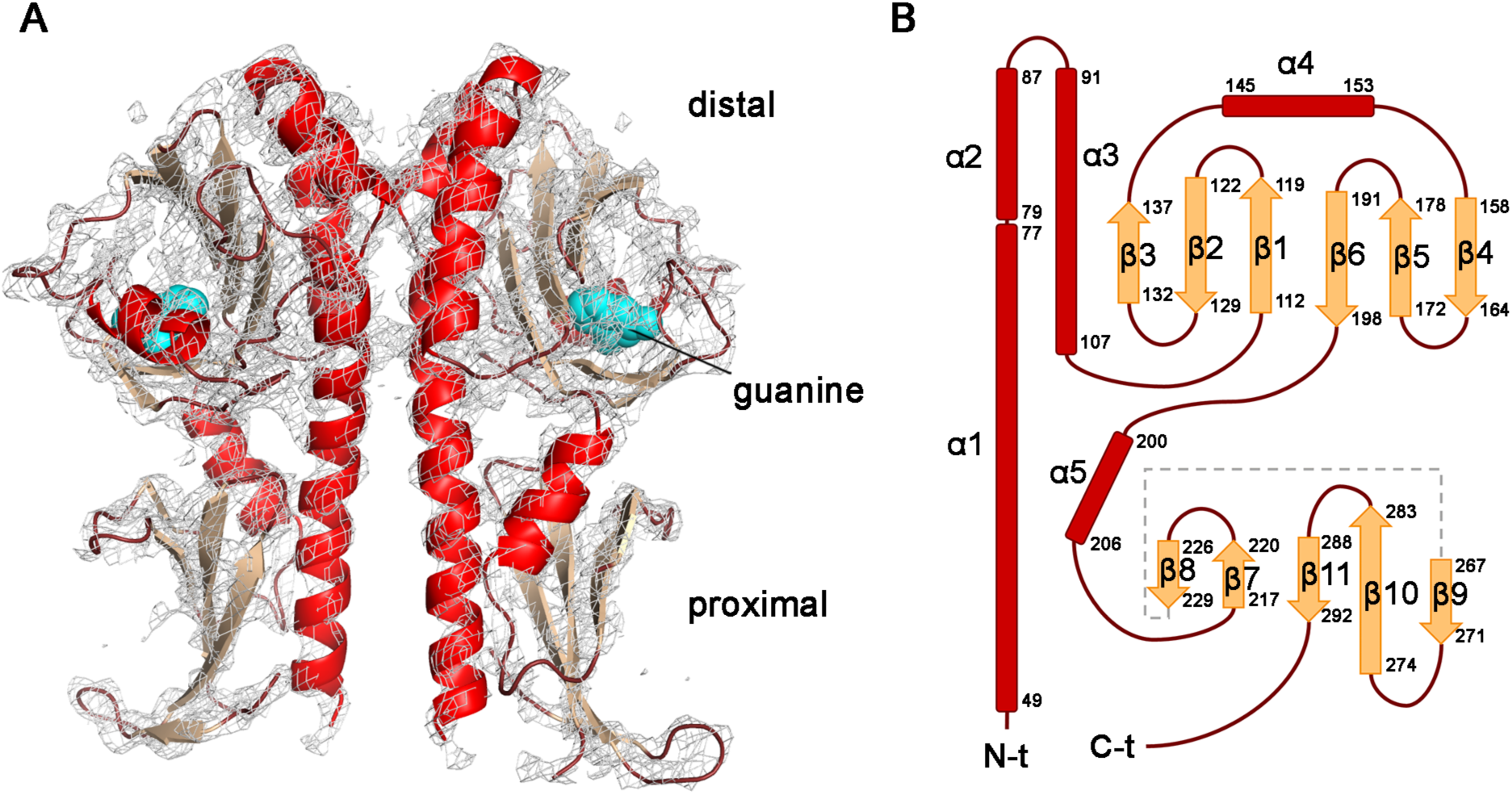
Tertiary and quaternary structure of Se-Met-Htc10-LBD. Similar results were obtained for Se-Met-Htc10-LBD_GUA_ and Se-Met-Htc10-LBD_HYP_. **(A)** The only Se-Met-Htc10-LBD molecule contained in the asymmetric unit forms a dimer with a nearby crystallographic symmetry mate, resembling the quaternary structure of other chemoreceptors LBDs; the 2*mFo- DFc* electron density map (gray mesh, 1.5 σ) evidences the ligand bound to the protein (cyan spheres). **(B)** A monomer is schematized, with residue numbers corresponding to full-length Htc10 provided for each secondary structure element. α-helices are shown in red and β-strands in orange.

*mFo-DFc* and 2*mFo-DFc* electron density maps evidenced the presence of the ligand (guanine or hypoxanthine) bound to the N-terminal domain of Se-Met-Htc10-LBD (Figure 2A). However, the low resolution of the structures of Se-Met-Htc10-LBD_GUA_ and Se-Met-Htc10-LBD_HYP_ did not allow determining details about the established interactions. In order to gain further insights about ligand binding to Htc10-LBD, we solved a high-resolution crystal structure of the native protein in complex with guanine.

### A high-resolution crystal structure of Htc10-LBD in complex with guanine

Crystallization assays of native Htc10-LBD plus guanine (Htc10-LBD_GUA_) resulted in crystals belonging to the space group P2_1_2_1_2_1_ that diffracted to high resolution (Table 2). The crystal structure contains six protein molecules in the asymmetric unit, encompassing Htc10 residues 49-208, while no electron density was evident in *mFo-DFc* or 2*mFo-DFc* maps for the most N-terminal residues or the C-terminal domain of the protein. The mean RMSD between Htc10-LBD chains is 0.8 for 154 aligned residues. Htc10-LBD monomers form three equivalent dimers with C2 symmetry, adopting a quaternary arrangement comparable to that of Se-Met-Htc10-LBD (Figure 3A-B). The interface area in Htc10-LBD dimers is *ca.* 800 Å^2^ and, similarly to Se-Met- Htc10-LBD, the dimeric interaction is stabilized by residues located in segment 70-107. The RMSD between Htc10-LBD and Se-Met-Htc10-LBD chains is around 1 Å for 142 aligned residues. These observations suggest that, at least in the presence of the ligand, the 3D architecture of Htc10-LBD and its intermolecular interactions are independent of the C-terminal domain of the protein. It is unclear whether the C-terminal domain of Htc10-LBD is highly flexible or it was degraded during crystallization: according to estimates of the Matthews coefficient, the presence of six whole molecules of Htc10-LBD in the asymmetric unit would imply a relatively low solvent content (30%); on the other hand, Htc10-LBD crystals grew after 45 days and the PeptideCutter server (https://web.expasy.org/peptidecutter/) predicts several protease cleavage sites near the most C-terminal segment of Htc10-LBD evidenced in electron density maps.

**Figure 3.**
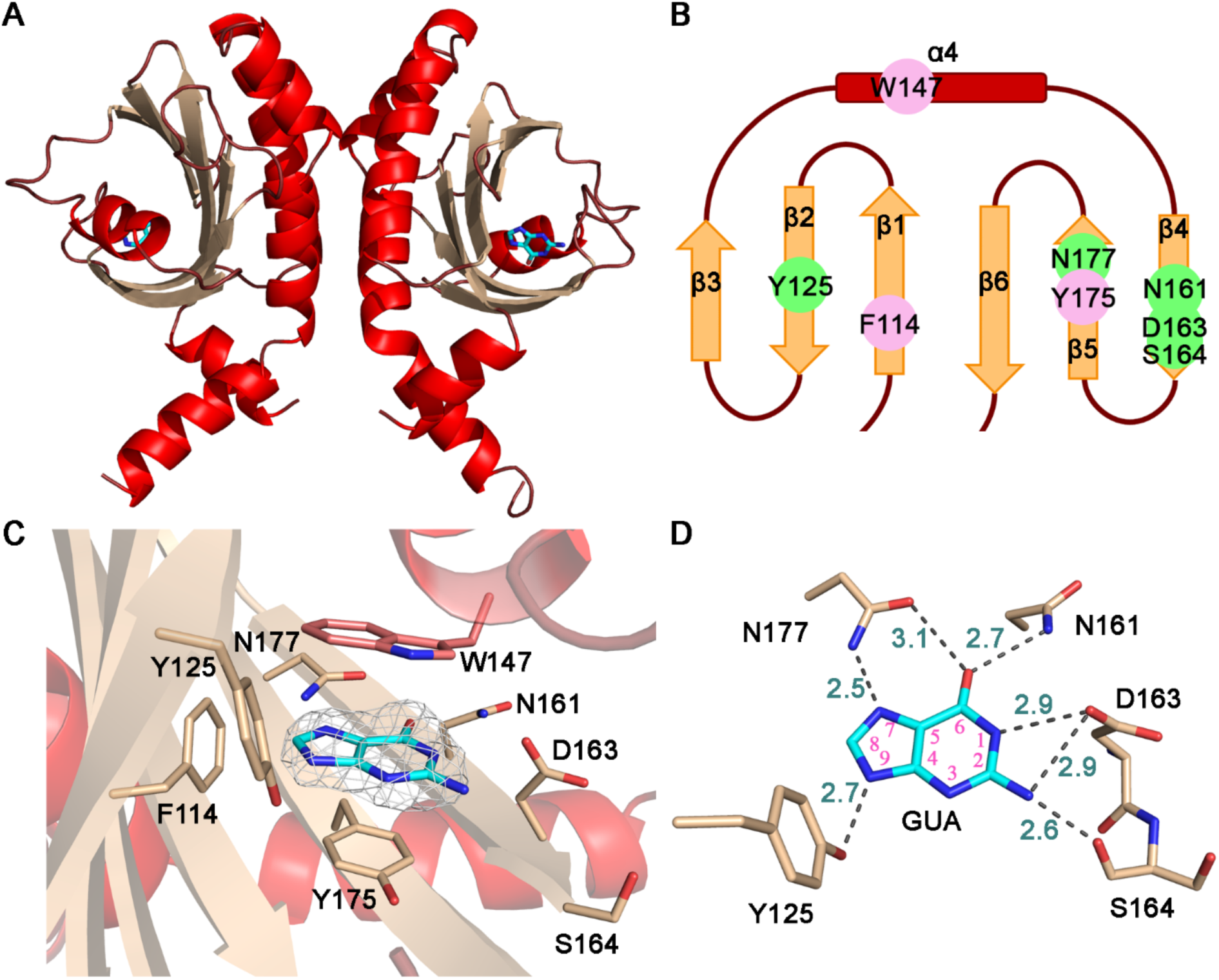
A high-resolution crystal structure of Htc10-LBD_GUA_. **(A)** Native Htc10-LBD forms a dimer with the same symmetry as Se-Met-Htc10-LBD (ribbons). α-helices are shown in red and β-strands in orange. **(B)** Schematic representation of the membrane distal domain showing the position of the hydrophobic (pink) and hydrogen bonding (green) residues that conform the ligand binding pocket. **(C)** Purine binding site. The 2*mFo-DFc* electron density map (gray mesh, 1.5 σ) is shown for the guanine molecule. The side chains of residues providing hydrophobic interactions (F114, W147 and Y175) as well as those of residues involved in hydrogen bonds with the ligand (Y125, N161, D163, S164 and N177) are shown in stick representation. **(D)** Hydrogen bonds links established between the guanine molecule and Htc10-LBD. Interatomic distances (green numbers) are informed in Å and depicted as dashed lines. Pink numbers indicate the position of atoms in the aromatic rings of guanine.

*mFo-DFc* and 2*mFo-DFc* electron density maps clearly evidenced a guanine molecule bound to each Htc10-LBD chain (Figure 3C-D). The protein-ligand hydrogen bond interactions revealed by the crystal structure of the Htc10-LBD_GUA_ complex involve residues Y125, N161, D163, N177 (side chains) and S164 (main chain). Besides, ligand binding is also stabilized by the side chains of W147 and Y175, which lie in parallel planes less than 4 Å above and below the guanine molecule, providing an hydrophobic environment. The side chain of F114 also establishes hydrophobic interactions with the ligand.

### Amino acids involved in ligand-binding

To understand the contribution of the predicted hydrogen bond interactions on ligand binding stabilization, we performed site directed mutagenesis targeted to the residues identified above. The amino acid replacements Y125F, N161A, D163A and N177A were introduced in different combinations and Htc10-LBD variants were produced and purified to perform ligand-binding experiments. In ligand-free thermal shift assays, all variants showed a decrease in *T*_m_ as compared to the wild-type protein, suggesting that the amino acid replacements affected the stability of the domain (Table 3). However, ligand-binding stabilization was affected as well. The variant with all four substitutions (Htc10-LBD-**FAAA**) became seemingly unable to bind guanine or hypoxanthine, as its *T*_m_ was unaffected by the presence of any of these ligands. The other two variants still showed some stabilization by the presence of the ligands, but the magnitude of the *T*_m_ shift was clearly reduced (Table 3). The corresponding dissociation constants were determined by isothermal calorimetry (Table 3 and Figure S1). For the native LBD, *K*_d_ values of 7 μM for guanine and 2.5 μM for hypoxanthine were determined (Table 3). These values were comparable to the ones estimated by thermal shift assays (see above), further validating the thermal shift-based approach for estimation purposes. The removal of the OH group from Y125 in Htc10-LBD Y125F (Htc10-LBD-**F**NDN) caused a decrease in the affinity for both ligands and a further reduction was driven by the additional substitution of both N161 and D163 (Htc10-LBD- **FAA**N). In all, thermal shift and isothermal titration calorimetry analyses support the involvement of these four residues in ligand binding.

**Table 3.** Effect of amino acid replacements in Htc10-LBD on the binding of guanine and hypoxanthine. Thermodynamic parameters were determined for Htc10-LBD and three variants with amino acid replacements at positions Y125, N161, D163 and N177 by isothermal titration calorimetry (ITC). *T*_m_ was determined using the thermal shift assay in the absence of ligands or in the presence of hypoxanthine (500 µM) or guanine (600 µM). Replaced amino acids are in bold. Gua: guanine; hyp: hypoxanthine; ND: not determined.

| LBD variant | $T_m$ (°C) | $\Delta T_m$ (°C) | | $K_d$ (ITC, μM) | | $\Delta H$ (kJ/mol) | | $\Delta S$ (J/K·mol) | | $\Delta G$ (kJ/mol) | |
| --- | --- | --- | --- | --- | --- | --- | --- | --- | --- | --- | --- |
|  |  | gua | hyp | gua | hyp | gua | hyp | gua | hyp | gua | hyp |
| Htc10-LBD-YNDN | 54.2<br>$\pm 0.2$ | 5.2<br>$\pm 0.1$ | 10.6<br>$\pm 0.2$ | 7.0<br>$\pm 0.6$ | 2.5<br>$\pm 0.3$ | -80<br>$\pm 9$ | -260<br>$\pm 20$ | -180<br>$\pm 20$ | -770<br>$\pm 80$ | -29.4<br>$\pm 0.8$ | -32.0<br>$\pm 0.9$ |
| Htc10-LBD-FNDN | 47.0<br>$\pm 0.2$ | 2.6<br>$\pm 0.2$ | 9.6<br>$\pm 0.2$ | 25<br>$\pm 2$ | 13.9<br>$\pm 0.9$ | -690<br>$\pm 40$ | -100<br>$\pm 10$ | -2200<br>$\pm 300$ | -240<br>$\pm 20$ | -26.3<br>$\pm 0.9$ | -27.7<br>$\pm 0.6$ |
| Htc10-LBD-FAAN | 45.1<br>$\pm 0.4$ | 3.0<br>$\pm 0.1$ | 4.9<br>$\pm 0.1$ | 118<br>$\pm 9$ | 140<br>$\pm 9$ | -180<br>$\pm 20$ | -550<br>$\pm 60$ | -540<br>$\pm 40$ | -1760<br>$\pm 90$ | -22.4<br>$\pm 0.7$ | -22<br>$\pm 1$ |
| Htc10-LBD-FAAA | 50.9<br>$\pm 0.6$ | 0.5<br>$\pm 0.4$ | 0.2<br>$\pm 0.2$ | ND | ND | ND | ND | ND | ND | ND | ND |

### A purine-chemotaxis receptor from *Pseudomonas putida* shares the Htc10 ligand-binding residues

Thermal shift assays have been extensively used to find out the specificity of a huge variety of receptors that mediate chemotaxis [18, 19]. One of the 27 chemoreceptors from *Pseudomonas putida*, McpH, was found to be the only chemoreceptor that mediates chemotaxis to purine derivatives such as adenine, guanine, xanthine, hypoxanthine and uric acid in that organism [20]. We wondered whether this chemoreceptor shared purine-binding determinants with Htc10. McpH displays a ligand-binding domain with a double-cache topology and 25% sequence identity with Htc10-LBD. An alignment between both LBDs shows a remarkable conservation of the amino acids found to be involved in purine binding in Htc10 (Figure 4A). Four out of seven residues whose side chains participate in ligand binding are identical (Y125, W147, N161 and D163 in Htc10), whereas the remaining three display conservative replacements (F114, Y175, N177). Moreover, the superposition of the crystallographic model of Htc10-LBD and that of McpH-LBD predicted by Alpha Fold [21, 22] shows a very good correspondence between the secondary structure elements that conform the membrane distal domain (Figure 4B), as well as a similar arrangement of residues involved in ligand binding (Figure 4C), suggesting that the identified residues participate in purine binding also in McpH. According to the crystal structure of McpH reported in a recent preprint, such assumption is confirmed [23]. Moreover, a consensus sequence is proposed for the prediction of purine binding ability in other double- cache domains. However, the suggested consensus sequence does not completely fit what we observed for Htc10, as it would identify R136 (R129 in McpH) as being involved in ligand binding, but such a residue is not part of the Htc10 binding pocket. Careful analyses of more purine-binding domains will help to refine the consensus sequence.

**Figure 4.**
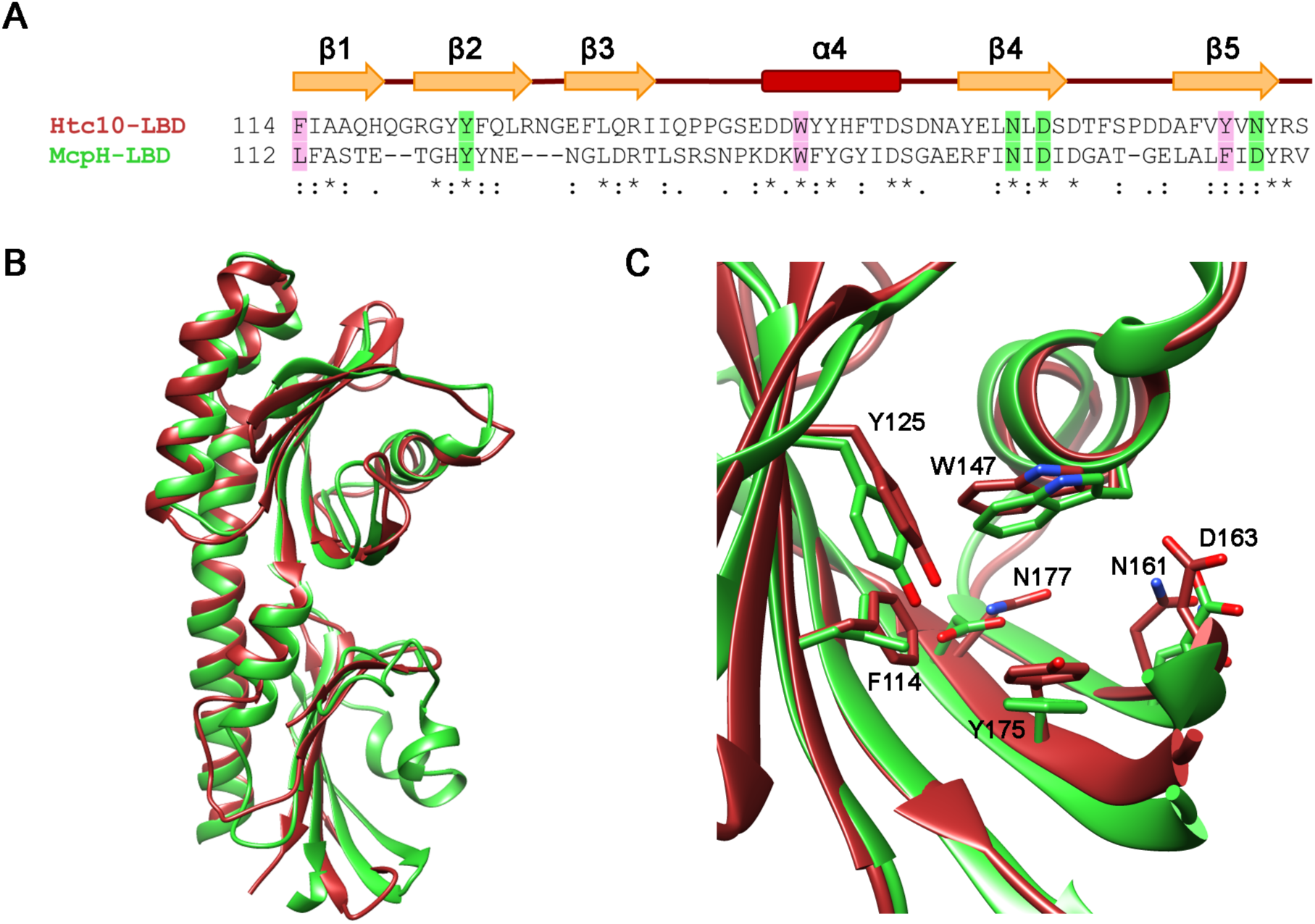
Comparison of Htc10-LBD and McpH-LBD. **A.** Sequence alignment between the two LBDs. * indicates identical conserved residues; : signals conservation between groups of strongly similar properties; . indicates residues with weakly similar properties; no symbol means that residues are not conserved. Residues that provide hydrophobic interactions with the ligand are highlighted in pink while those involved in hydrogen bonding are shown in green. A schematic representation of the secondary structure is shown on top of the alignment. **B.** Superposition of the crystallographic model of Htc10-LBD (red ribbon) and the Alpha Fold model of McpH-LBD (green ribbon). **C.** The ligand-binding pocket. Residues highlighted in the sequence alignment are shown in stick representation.

Although the possibility of predicting the specificity of chemoreceptors based solely on their sequence and/or folding type is incipient, the accumulation of information keeps opening ways toward this end. For instance, consensus sequences for double-cache domains that bind amino acids have been described which have prediction ability [24, 25]. Also, a consensus sequence has been proposed for biogenic amines [26].

### Htc10-like chemoreceptors are broadly distributed and show a highly conserved cytoplasmic domain

The chemotaxis-related *Ht*Che2 cluster found in the genome of *Halomonas titanicae* KHS3 was initially described as a putative Wsp-like signal transduction pathway due to the presence of a gene coding for a protein comprising a diguanylate cyclase domain and phosphorylatable receiver domains. Its enzymatic activity could thus be presumably controlled to modulate c-di-GMP production [3]. However, many features differentiate both pathways, not only in terms of gene content, but also in the order of the genes and their domain composition. Notably, a HMMER [27] search of the *Ht*Che2 cluster revealed similar gene clusters in distant bacteria, including different orders and families within the alpha, beta, and gamma Proteobacteria [3].

Therefore, we wondered whether the chemoreceptor controlling such a pathway was conserved. An alignment of Htc10-like receptors found in fourteen different *Ht*Che2-like gene clusters evidenced a relatively low conservation of the ligand-binding domain as compared to the cytoplasmic region of the protein. Only those LBDs with the highest sequence identity (36- 38%) (Table 4) with Htc10-LBD, found in *Nitrincola* and *Marinospirillum* genera within the Oceanospirillales order, conserve the predicted double-cache fold as well as residues involved in purine binding (Figure 5A), suggesting that in these cases the receptors might share specificity. While the average sequence identity along the full-length of the analyzed Htc10-like receptors reached 35% (Table 4), in most cases it was less than 20% for the LBD. Highlighting such diversity, three LBDs would not even adopt a double-cache fold.

**Figure 5.**
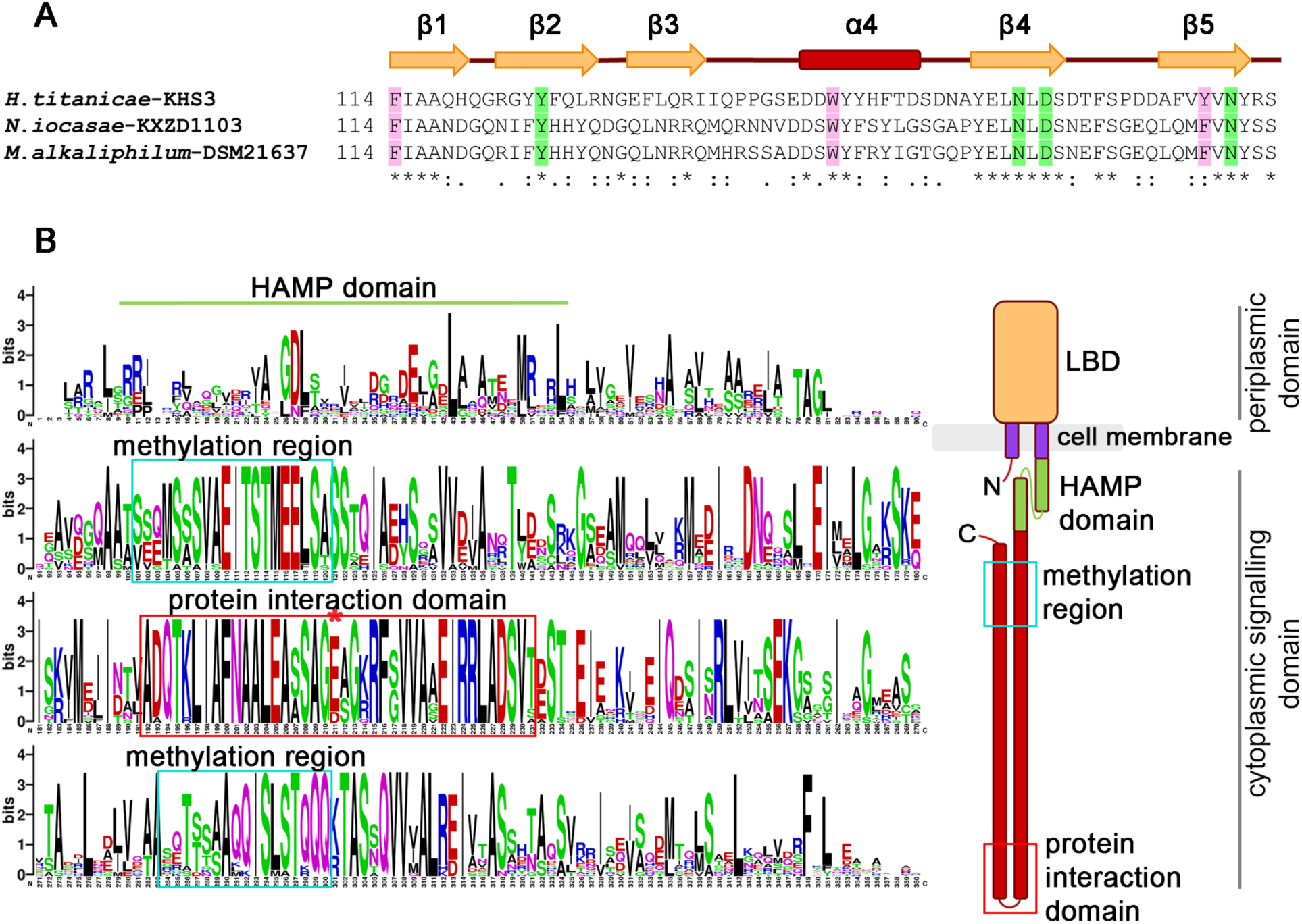
Htc10-like chemoreceptors from other microorganisms. **A.** Comparison of the LBDs of Htc10-like chemoreceptors belonging to the order Oceanospirillales, which includes *Halomonas*. Residues that provide hydrophobic interactions with the ligand are highlighted in pink while those involved in hydrogen bonding are shown in green. A schematic representation showing the secondary structure elements of the corresponding sequences is shown at the top. **B.** Weblogo of the Htc10-like cytoplasmic domain. Fourteen sequences are compared, each from one genus in which the *Ht*Che2-like cluster was found (species detailed in Table 4). The region of the HAMP domain is indicated, as well as the two sequences that correspond to the methylation region (light blue rectangles), and the region corresponding to the highly conserved hairpin tip that is expected to be involved in protein interactions (red rectangle). The Glu residue that is at the very tip of the cytoplasmic hairpin is marked with a red asterisk. On the right, a schematic representation of Htc10 is shown, in which the methylation region and the protein interaction region are indicated.

**Table 4.** Comparison of Htc10 with other chemoreceptors. The first part of the table shows the sequence identity between Htc10 and chemoreceptors coded within highly similar *Ht*Che2- like clusters identified in other Proteobacteria [3]. Pairwise identities (%) are provided for full- length proteins, periplasmic domains and cytoplasmic domains. The folding type predicted by Swiss-Model (https://swissmodel.expasy.org) for the LBDs is stated between parentheses. A similar comparison is shown below for McpH and WspA.

|  | Order | Genus | Full-length protein | Periplasmic domain | Cytoplasmic domain |
| --- | --- | --- | --- | --- | --- |
| γ | Oceanospirillales | *Halomonas titanicae* KHS3 | 100 | 100 (dCache) | 100 |
|  |  | *Marinospirillum alkaliphilum* DSM21637 | 55 | 38 (dCache) | 70 |
|  |  | *Nitrincola iocasae* KXZD1103 | 53 | 36 (dCache) | 67 |
|  | Chromatiales | *Thiorhodospira sibirica* A12 | 41 | 11 (4HB-like) | 56 |
|  |  | *Thiorhodovibrio* sp. 970 | 38 | 13 (4HB-like) | 50 |
|  |  | *Allochromatium vinosum* DSM180 | 36 | 18 (dCache) | 51 |
|  |  | *Thiorhodococcus drewsii* AZ1 | 36 | 17 (dCache) | 50 |
|  |  | *Thiocapsa marina* 5811 | 36 | 19 (dCache) | 49 |
|  |  | *Marichromatium purpuratum* 984 | 36 | 17 (dCache) | 50 |
|  | Enterobacteriales | *Buttiauxella agrestis* CDC 1176-81 | 36 | 10 (dCache) | 51 |
|  |  | *Citrobacter youngae* ATCC 29220 | 35 | 14 (dCache) | 50 |
| α | Magnetococcales | *Magnetococcus marinus* MC-1 | 38 | 17 (dCache) | 52 |
|  | Rhodospirillales | *Pararhodospirillum photometricum* DSM122 | 37 | 17 (dCache) | 52 |
|  |  | *Magnetospirillum magneticum* AMB-1 | 37 | 17 (dCache) | 52 |
| β | Rhodocyclales | *Thauera chlorobenzoica* 3CB1 | 35 | 14 (4HB) | 45 |
| McpH | *Pseudomonas putida* Chemotaxis receptor |  | 27 | 25 (dCache) | 30 |
| WspA | *Pseudomonas aeruginosa* Wsp pathway chemoreceptor |  | 23 | 22 (4HB) | 25 |

In contrast, the sequence identity for the cytoplasmic domain of the analyzed Htc10-like receptors was near or higher than 50% (Table 4). Such figure raised to 95-100% in a number of selected positions located at the tip of the predicted cytoplasmic hairpin and at the methylation region (Figure 5B). This scenario resembles that of chemotaxis-controlling receptors, where a highly conserved cytoplasmic domain that interacts with other chemoreceptors and downstream signalling proteins can present very diverse sensor domains, depending on the variety of stimuli to which the pathway needs to respond and reflecting differences in inhabited niches and/or lifestyles of the microorganisms. Likewise, a recent study on the chemoreceptor present in almost 800 gene clusters closely related to the Wsp pathway of *P. aeruginosa* showed a broad diversification of the sensor domain [28]. The same study showed that such pathways were largely restricted to the genus *Pseudomonas* within gamma Proteobacteria and to some representatives of beta Proteobacteria.

When Htc10 was compared to WspA, the chemoreceptor that controls the Wsp pathway in *P. aeruginosa*, the sequence identity between the full-length proteins as well as between the periplasmic or cytoplasmic portions was around 20-25%, without an apparently higher similarity for any of the domains (Table 4). The low identity value between the periplasmic domains of these two receptors was expected, as their LBDs do not share folding type. However, big differences between their cytoplasmic domains are rather surprising taking into account that *Ht*Che2 pathway seems to be functionally equivalent to the Wsp [4]. The fact that some other components show important differences between pathways (for example regarding coupling proteins and domains in the associated diguanylated cyclase) highlights the need of more studies to characterize the signalling behavior of the proteins involved in these cases and also in related pathways in other microorganisms.

The main physiological role of the *Ht*Che2 transduction pathway could be still considered an open question. Even though the removal of the cognate methyltransferase gene and the consequent increase in receptor methylation generates an hyperbiofilm phenotype that is comparable to the equivalent mutant in the *Pseudomonas* Wsp pathway [4], it is likely that this only reflects an increase in the activity of the controlled diguanylate cyclase and not necessarily reveals the main function of the pathway. The strong conservation of the Htc10-like cytoplasmic domain, its divergence compared to WspA-like receptors as well as additional differences in gene and/or domain composition of the predicted signalling components, leave open the question about how related both pathways are in terms of signalling. However, the fact that both pathways are expected to control the activity of an associated diguanylate cyclase might allow extrapolations and highlights the importance of ligands that can modulate the activity of the enzyme via their binding to the corresponding chemoreceptors.

### Concluding remarks

We have characterized the ligand-binding domain of Htc10, the chemoreceptor controlling the *Ht*Che2 pathway in the marine bacterium *Halomonas titanicae* KHS3. Htc10-LBD binds the purine derivatives hypoxanthine and guanine with high affinity, with dissociation constants in the micromolar range. Residues involved in ligand binding were identified, both those that provide a hydrophobic environment for the nitrogenous base as well as those involved in specific hydrogen bonds with the ligand. Substitution of the latter resulted in a protein unable to bind ligands.

The strong conservation of Htc10-like chemoreceptors and their distribution among distant proteobacterial lineages suggests that the pathway has been actively incorporated and/or maintained under selection pressure, and might play a significant role in the organisms that harbor it. The identification of specific ligands for one of such receptors, presumably controlling the activity of an associated diguanylate cyclase, opens the way to studies focused on the dynamics of kinase activation in response to such compounds. Also, it will be possible to study whether this receptor displays an adaptation response similar to the one observed for the chemotaxis receptors. This kind of signalling might have similarities with the WspA-like chemoreceptors that have periplasmic domains for up to now unidentified ligands. Further studies will be needed to unveil the physiological role of the *Ht*Che2 pathway, and its relationship with sensing of purine derivatives.

## Materials and Methods

### Cloning

The sequence encoding the periplasmic domain of the Htc10 chemoreceptor from *Halomonas titanicae* KHS3, Htc10-LBD (94-927), was amplified by a polymerase chain reaction (PCR) using purified genomic DNA as template and primers nFR3 and nFR8 (Table S1). The reaction mixture contained 1.5 U Taq (PB-L Biological Products), 0.5 mM dNTPs (PB-L Biological Products), 3 mM MgCl_2_ (PB-L Biological Products), 0.25 µM of each oligonucleotide and 1× reaction buffer (PB-L Biological Products). The PCR cycles were as follows: (95 °C, 1 min) × 1; (95 °C, 30 s; 50 °C, 30 s; 72 °C, 1 min) × 31; (72 °C, 5 min) × 1, in a T100TM Thermal Cycler (Bio-Rad). The product was visualized by electrophoresis on 1% w/v agarose gels in 1× TBE buffer. The amplified product (previously purified using EasyPure Quick Gel Extraction Kit, Transgenbiotech) and the pET28a expression vector (obtained with the EasyPure Plasmid MiniPrep Kit) were digested with the NcoI (Thermo Fisher) and XhoI (Thermo Fisher) restriction enzymes. Reactions were carried out under the conditions recommended by the manufacturer. Ligation was performed using T4 DNA ligase (Promega) with 3:1 insert:vector ratio in the buffer provided by the manufacturer. Incubation was carried out overnight at 4 °C.

For preparation of competent cells, 25 mL of LB were inoculated with 1% v/v of a saturated *E*. *coli* culture and incubated at 37 °C. After 1.5 h cells were harvested (5000 × g for 10 min), resuspended in 10 mL of cold 0.1 M CaCl_2_ (Merck), and incubated for 30 minutes on ice. After this period, the cells were centrifuged again, and finally they were resuspended in 2.5 mL of 0.1 M CaCl_2_ (at 4 °C) with 15% v/v glycerol. 200 µL of competent cells were incubated with the ligation product on ice for 30 minutes. After a 2-minute heat shock at 42 °C, 800 µL of LB were added and the cells were incubated for 30 minutes at 37 °C. Finally, transformed cells were selected on LB plates supplemented with 50 µg/ml kanamycin. The presence of the Htc10-LBD gene was verified by PCR and confirmed by sequencing (Macrogen Inc, Korea). The obtained construct was named pFR2 (Table S2).

### Site-directed mutagenesis

Mutations in Htc10-LBD were generated through site-directed mutagenesis by PCR (Quick Change) using plasmid pFR2 as template and DNA polymerase Pfx (Thermo Fisher). Oligonucleotides carrying the desired mutations are listed in Table S1. The PCR program was: (94 °C, 1 min) × 1; (94 °C, 1 min; 58 °C, 1 min; 68 °C, 15 min) × 21; (68 °C, 10 min) × 1. The amplification product was digested with DpnI (Thermo Fisher) for 1.5 h at 37 °C and then used to transform competent *E. coli* Top10 cells. The constructs were confirmed by sequencing of the coding region.

### Production of pure recombinant Htc10-LBD

Optimal conditions for Htc10-LBD expression were determined in *E. coli* BL21 cells transformed with plasmids pFR2 and pKJE7, in LB medium supplemented with 50 µg/mL kanamycin and 25 µg/mL chloramphenicol. Protein production was induced with 0.1 or 0.5 mM isopropyl-β-D-1- thiogalactopyranoside (IPTG, PB-L Productos Bio-lógicos), with or without 0.5% w/v L-arabinose (Merck), with agitation for 4 hours or overnight, at 18, 28 or 37 °C. Subsequently, the culture was centrifuged at 5000 × g for 10 minutes at 4 °C, and the pellet was resuspended to optical density 20 with 25 mM Tris-HCl, pH 8.0, supplemented with 300 mM NaCl, 1 mg/mL lysozyme from egg white (Genbiotech), and 1 mM phenylmethylsulfonyl fluoride (PMSF, Sigma Aldrich). This suspension was sonicated using a Sonics Vibra-Cell VCX-600 Ultrasonic Processor, with 60 cycles of 2 s sonication and 3 s rest. The resulting extract (total proteins) was centrifuged for 45 minutes at 23000 × g at 4 °C, and then both the pellet (insoluble fraction) and the supernatant (soluble fraction) were separated and stored at -20 °C. In subsequent experiments, the soluble fraction was obtained from 1 L of culture, induced overnight with 0.1 mM IPTG at 18 °C, following the protocol just described.

SeMet-Htc10-LBD was expressed in *E. coli* BL21(DE3) transformed with plasmids pFR2 and pJEK7, in media supplemented with 50 µg/mL kanamycin and 25 µg/mL chloramphenicol. Transformed cells grown for 5.5 h at 37 °C in 2YT medium were used to inoculate M9 minimal medium at 1:100 and kept overnight at 37 C° with constant shaking. The overnight culture, showing an OD_600_ ∼ 2.3, was then used to inoculate 5 L of fresh M9 medium (1:50), grown at 37°C until OD_600_ reached 0.6, when the biosynthesis of methionine was inhibited by adding lysine, phenylalanine, and threonine at 100 mg/L, isoleucine and valine at 50 mg/L, and providing selenomethionine at 60 mg/L. 30 min following the supplementation with the amino acids listed above, SeMet-Htc10-LBD expression was induced by adding IPTG to a final concentration of 0.1 mM, and leaving the culture at 18 °C for 30 h with constant shaking. Cell pellets were harvested and frozen at -80 °C until protein purification which was performed.

In all cases, Htc10-LBD (or SeMet-Htc10-LBD) production was analyzed by SDS-PAGE. Htc10-LBD was purified using immobilized metal affinity chromatography (IMAC) on ÄKTA Explorer 100 (GE Healthcare) at 4 °C. The soluble crude protein extract was loaded onto HisTrap 1 mL columns, previously equilibrated with 25 mM Tris-HCl pH 8.0, 300 mM NaCl, 10 mM imidazole (Merck). Elution was performed in the same buffer but with a linear gradient of imidazole up to 300 mM. Fractions were collected in 2 mL aliquots while monitoring absorbance at 280 nm and examined by SDS-PAGE. The most concentrated protein fractions were pooled and quantified.

When protein samples were intended for crystallization, Htc10-LBD underwent additional purification by size-exclusion chromatography on a preparative scale. For this, the fractions obtained by affinity chromatography were pooled and concentrated using conical membrane ultrafiltration tubes (Vivaspin 6 mL 10 kDa MWCO, Sartorius) to 7 mg/mL (2 mL). The resulting sample was loaded onto a Superdex 200 column (GE Healthcare) and elution was performed with 25 mM Tris-HCl pH 8.0, 300 mM NaCl. Fractions from this chromatography were collected in 5 mL aliquots, evaluated by SDS-PAGE, concentrated by ultrafiltration, and stored at -80 °C for subsequent crystallization. Protein concentration was determined by absorbance at 280 nm or using the Bradford method [29]. SeMet-Htc10-LBD purification was performed following the same protocol as for native Htc10-LBD.

### Thermal Shift Assay

Incubation mixtures (30 µl) contained 10 µM of protein and 5× SYPRO Orange (Sigma) in 25 mM Tris-HCl pH 8.0, 300 mM NaCl, in the absence of any ligand or in the presence of the tested compounds. Biolog plates (Hayward, CA, USA) screening was done using mixtures containing 10 µl of each resuspended compound, rendering an estimated final concentration between 3 and 6 mM. Selected compounds were individually re-tested at 10 mM final concentration. Concentration curves were done with guanine (0-1200 µM, Sigma Aldrich) or hypoxanthine (0-10000 µM, Sigma Aldrich). Incubation mixtures were arranged in 96-well plates and introduced into the StepOne Real-Time PCR equipment (Thermo Fisher Scientific). Samples were heated from 10 °C to 80 °C and fluorescence was measured at each point with SYBR Green filters. The melting temperature (*T*_m_) was determined as the minimum value in the first derivative curve. All thermal shift assays performed with known ligand concentration were carried out in triplicate. The observed dispersion between estimated *T*_m_ values did not exceed 5% in different experiments.

### Isothermal Titration Calorimetry

ITC analyses were performed at the facility of Instituto de Biología y Medicina Experimental (IBYME), Buenos Aires, Argentina. Measurements were carried out using a NanoITC calorimeter (TA Instruments) at 25 °C. Hypoxanthine and guanine solutions (0.25-1 mM) were prepared in 25 mM Tris-HCl pH 8.0, 300 mM NaCl. Protein samples (25-70 µM) were titrated with 20 successive injections of 2.5 µL ligand solution at a spacing of 300 s. The titration cell was continuously stirred at 300 rpm. The heats of dilution of the ligands in the buffer were subtracted from the titration data. Data analysis was performed using Nano Analyze Software.

### Htc10 crystallization, data collection and structure determination

Crystallization trials of seleno-methionine labeled Htc10-LBD (Se-Met-Htc10-LBD, 32 mg/ml) plus guanine or hypoxanthine (2 mM in both cases) were carried out using the sitting-drop vapor diffusion method as performed at the Crystallography Platform of the Institut Pasteur, according to established protocols [30]. Crystallization trials of native Htc10-LBD (10 mg/ml) plus guanine (5 mM) were also carried out using the sitting-drop vapor diffusion method but employing a Gryphon nanoliter-dispensing crystallization robot (Art Robbins Instruments), by mixing 250 nl of protein solution and 250 nl of reservoir solution, equilibrated against 80 µl of reservoir solution, in 96-well Intelli-Plates (Hampton). In this case, crystallization plates were stored at 20°C and inspected with a stereo microscope (Leica) to monitor crystal growth.

The optimized conditions for crystal growth were 100 mM sodium acetate pH 4.6, 2 M sodium formate for Se-Met-Htc10-LBD plus guanine (Se-Met-Htc10-LBD_GUA_) crystals, 1 M (NH_4_)_2_SO_4_ for Se-Met-Htc10-LBD plus hypoxanthine (Se-Met-Htc10-LBD_HYP_) crystals and 200 mM (NH_4_)_2_SO_4_, 100 mM sodium cacodylate trihydrate pH 6.5, 30% w/v polyethylene glycol 8000 for Htc10-LBD plus guanine (Htc10-LBD_GUA_) crystals. Se-Met-Htc10-LBD_GUA_ and Se-Met-Htc10-LBD_HYP_ crystals grew after 12 and 5 days, respectively, whereas Htc10-LBD_GUA_ crystals grew after 45 days, to a size of *ca.* (50 µm)^3^ in all cases. Htc10-LBD_GUA_ crystals were flash-frozen in liquid nitrogen for data collection at 100 K. Se-Met-Htc10-LBD_GUA_ and Se-Met-Htc10-LBD_HYP_ crystals were treated similarly, using 25% v/v glycerol as cryo-protectant.

X-ray diffraction data from Se-Met-Htc10-LBD_GUA_ and Se-Met-Htc10-LBD_HYP_ crystals were measured on beamline ID30A-3 (European Synchrotron Radiation Facility, Grenoble, France) using wavelength 0.967697 Å, close to the peak of the Se K edge. On the other hand, X-ray diffraction data were collected from a single Htc10-LBD_GUA_ crystal on beamline I04 (Diamond Synchrotron, United Kingdom) using wavelength 0.9795 Å. In each case, the diffraction data were indexed and integrated with XDS [31] and scaled with Aimless [32] from the CCP4 program suite [33].

Se-Met-Htc10-LBD_GUA_ and Se-Met-Htc10-LBD_HYP_ crystals belonged to the space group H32 and diffracted to low resolution. The automated pipeline Crank-2 [34] was used with default parameters to determine the selenium substructure, calculate initial SAD phases, density modification and automatic model building for a Se-Met-Htc10-LBD_GUA_ crystal. The obtained electron density map was interpretable and allowed to recognize the topology of a double cache domain in the asymmetric unit, which together with a nearby crystallographic symmetry mate forms a protein dimer with C2 symmetry. However the automatically built model was incomplete and had poor stereochemistry. Therefore, an *ab initio* model of Htc10-LBD built using trRosetta [35] was rigid-body fitted into the electron density map and used to complete and correct the atomic coordinates of the polypeptide. The preliminary crystallographic model thus obtained was refined by iterative cycles of manual model building with Coot [36], employed to apply stereochemical restraints, and crystallographic refinement of atomic coordinates and individual B-factors using Refmac [37] with secondary structure restraints.

Native Htc10-LBD crystallized in the presence of guanine in space group P2_1_2_1_2_1_ and diffracted to high resolution. In this case, the crystal structure was solved by molecular replacement with Phaser [38] using as a search probe Htc10-LBD atomic coordinates available from the SAD experiment. Although searches performed with a single copy of the polypeptide failed, it was possible initially to place a C2 dimer in the asymmetric unit. After refining atomic coordinates and individual B-factors with Refmac [37], *mFo-DFc* and 2*mFo-DFc* maps showed clear evidence for the N-terminal portion of Htc10-LBD but instead the electron density vanished from residue 209 towards the C-terminus of each polypeptide. Therefore, such region was removed and the resulting model was further refined as previously described. Then, two more C2 Htc10- LBD dimers were placed in the asymmetric unit by molecular replacement. After successive cycles of manual model building with Coot [36] and crystallographic refinement with Refmac [37], the guanine molecules were manually placed in a *mFo–DFc* electron density map. The structure of Htc10-LBD_GUA_ was finally refined to convergence with phenix.refine [39].

The atomic coordinates of residues 66-208 in the preliminary structure of Se-Met-Htc10-LBD_GUA_ were replaced by the equivalents in a protein molecule in the crystallographic model of Htc10- LBD_GUA_. The structure was then refined by iterative cycles of manual model building with Coot [36] and crystallographic refinement of atomic coordinates and individual B-factors using Refmac [37] with external restraints generated from the Htc10-LBD model built by trRosetta [35]. Finally, the guanine molecule was manually placed in a *mFo–DFc* electron density map and the structure of Se-Met-Htc10-LBD_GUA_ was refined to convergence with phenix.refine [39]. The structure of Se-Met-Htc10-LBD_HYP_ was solved by molecular replacement with Phaser [38] using as a search probe the atomic coordinates of the polypeptide in the crystallographic model of Se-Met-Htc10-LBD_GUA_. The hypoxanthine molecule was manually placed in a *mFo–DFc* electron density map and the structure of Se-Met-Htc10-LBD_HYP_ was refined to convergence with phenix.refine [39].

The final models were validated through the Molprobity server [40].

Atomic coordinates and structure factors obtained for Se-Met-Htc10-LBD_GUA_, Se-Met-Htc10- LBD_HYP_ and Htc10-LBD_GUA_ were deposited in the Protein Data Bank under the accession codes 9B9S, 9B9X and 9BA3, respectively.

## Abbreviations

dNTP: Deoxyribonucleotide triphosphate
HAMP: <u>H</u>istidine kinase/<u>a</u>denylyl cyclase/<u>m</u>ethyl-accepting chemotaxis protein/<u>p</u>hosphatase
IPTG: Isopropyl thio-β-D-galactoside
LBD: Ligand-binding domain
MCP: Methyl-Accepting Chemotaxis Protein
SDS-PAGE: Denaturing polyacrylamide gel electrophoresis
WSP: Wrinkly Spreader Phenotype

## Author Contributions

Authors made substantial contributions in the following aspects: FERR, MKHS, MNL AND CAS: conceptualization, data acquisition/interpretation/analysis and writing; AFG, AB, KJ and MB: data acquisition/interpretation. María-Natalia Lisa and Claudia A. Studdert should be considered joint senior authors.

## Acknowledgements

This work was supported by research grants from the Agencia Nacional para la Promoción Científica y Tecnológica, Argentina, to CAS (PICT2016-1629; PICT2020-3814) and MNL (PICT-2019-00989) as well as by the Deutsche Forschungsgemeinschaft to KJ (JU270/19-1, project number 432402323). MKHS, MNL and CAS are career investigators from Consejo Nacional de Investigaciones Científicas y Técnicas (CONICET, Argentina). FERR and AFG held CONICET scholarships. AFG held also a DAAD-ALEARG fellowship. We thank Eduardo Bruch for his technical assistance in the crystallization of Se-Met-Htc10-LBD, the Crystallography Platform at the Institut Pasteur, namely C. Pissis and P. Weber, for performing robot-driven crystallization assays with SeMet-Htc10-LBD, and A. Haouz for helpful discussions. We also thank the synchrotron sources ESRF (Grenoble, France) and Diamond Light Source (Didcot, UK) for granting access to their facilities, and their respective staff for assistance in data collection.

## Supplementary material

**Table S1.**
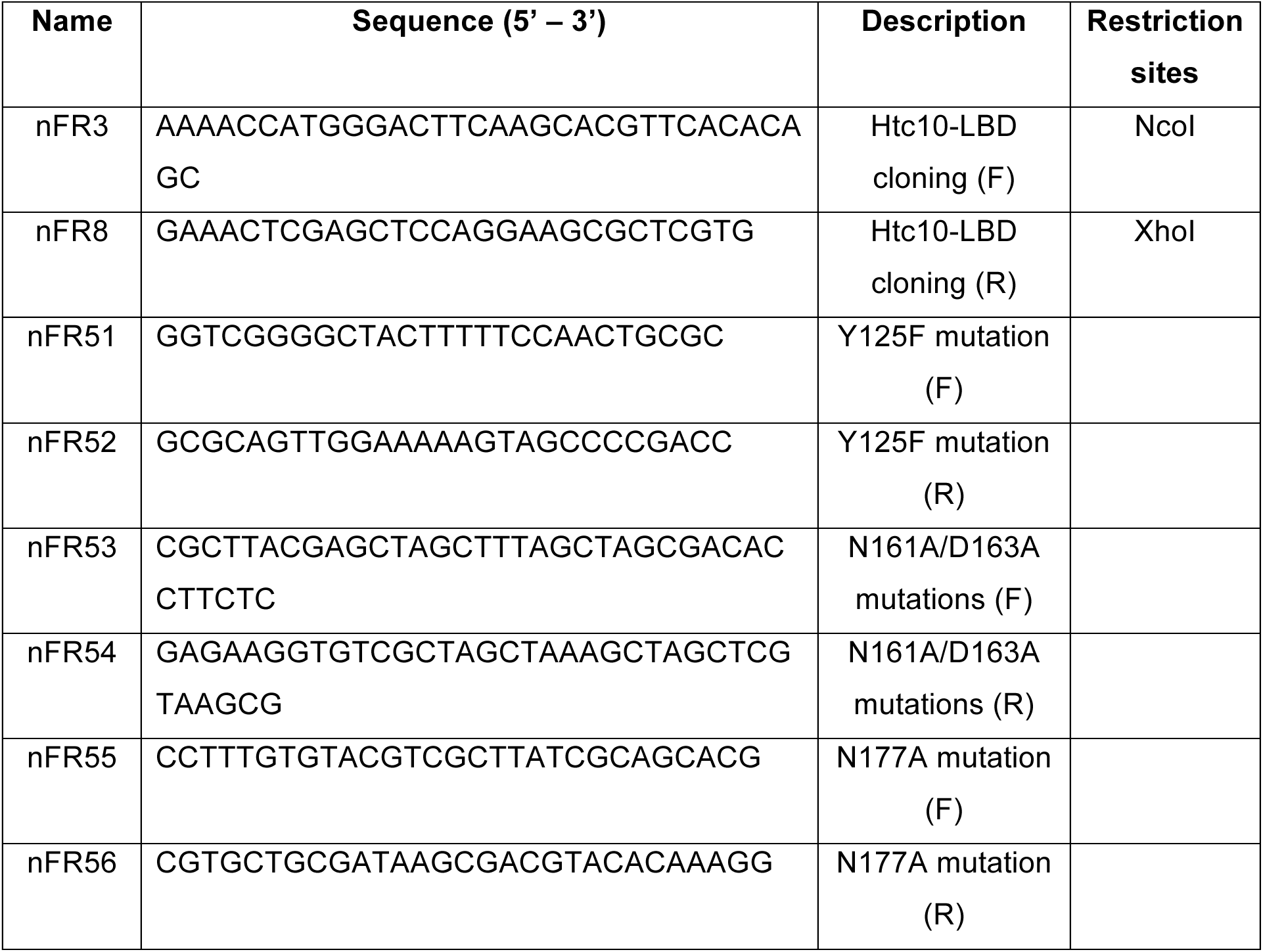
Oligonucleotides.

**Table S2.** Plasmids.

| Name | Characteristics | Source |
| --- | --- | --- |
| pET28a | KmR, IPTG-inducible expression vector | Novagen |
| pFR2 (YNDN) | pET28a derivative containing Htc10-LBD | This work |
| pHtc10-LBD-FNDN | pFR2 derivative containing Y125F mutation | This work |
| pHtc10-LBD-FAAN | pFR2 derivative containing Y125F, N161A and D163A mutations | This work |
| pHtc10-LBD-FAAA | pFR2 derivative containing Y125F, N161A, D163A and N177A mutations | This work |
| pKJE7 | CmR, DnaK-DnaJ-GrpE chaperone-coding L-arabinose-inducible | Schröder et al, 1993 |

**Figure S1.**
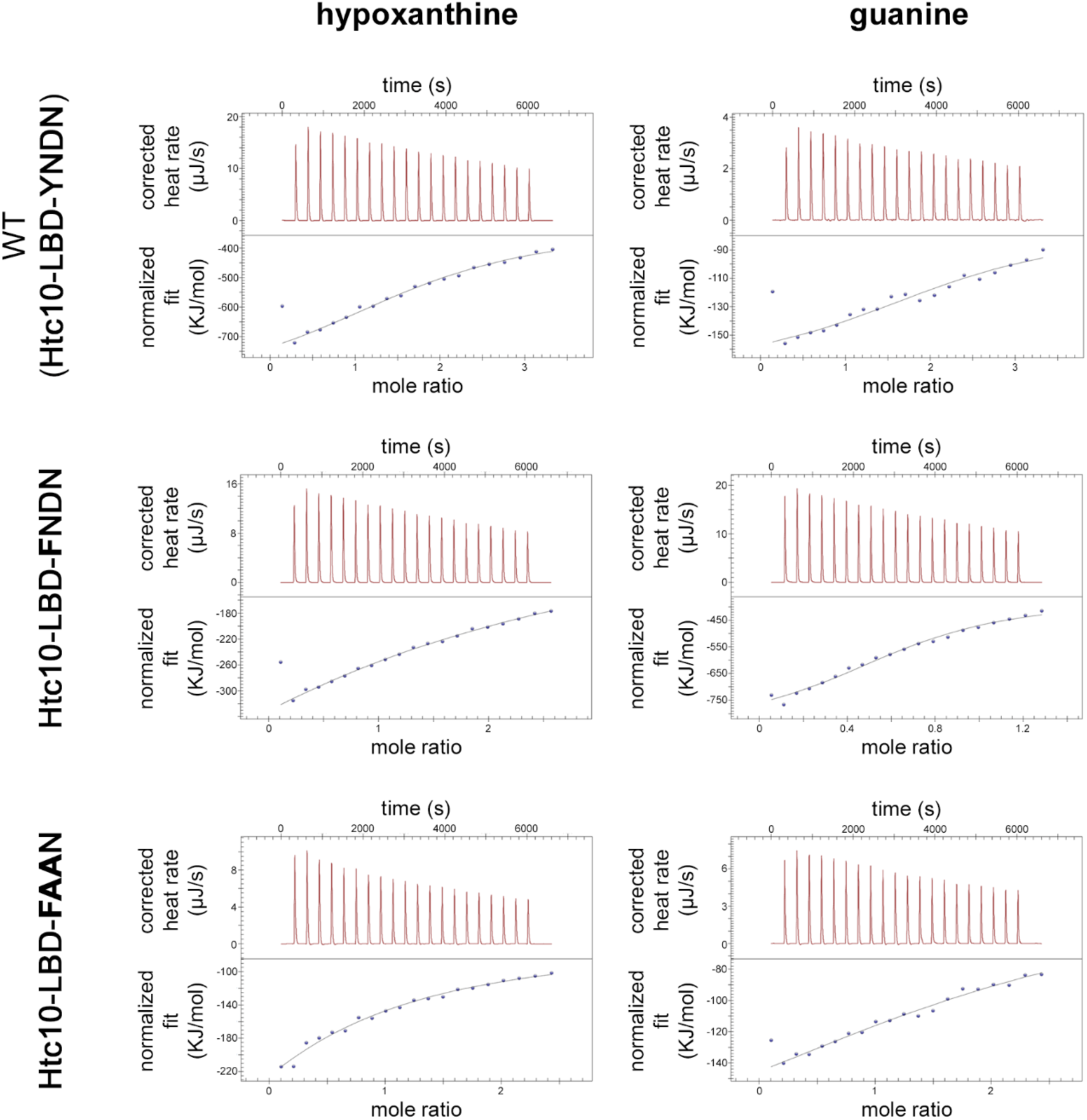
Raw data corresponding to the isothermal calorimetry analyses whose parameters are shown in Table 3. Thermograms of each protein obtained after the addition of guanine (right) or hypoxanthine (left). In each graph, raw heat release data per injection is shown (top) alongside heat normalized per ligand mole, along with the fitting curve (bottom).

